# A mouse model of myotonic dystrophy type 1 exhibits pain-like behavior and peripheral nociceptor hyperexcitability

**DOI:** 10.64898/2026.09.10.750655

**Authors:** Tyler S. Nelson, Aida Calderon-Rivera, Sara Hestehave, Santiago Loya-Lopez, Kimberly Gomez, Paz Duran, Rajesh Khanna

## Abstract

Pain is a prevalent and disabling symptom of myotonic dystrophy type 1 (DM1), yet its underlying mechanisms remain poorly understood. Using HSA LR20b transgenic mice, we found multimodal mechanical and thermal hypersensitivity. Whole-cell electrophysiological recordings demonstrated depolarized resting membrane potentials and increased action potential firing selectively in small- and medium-diameter dorsal root ganglion neurons. These findings identify peripheral nociceptor sensitization as a potential mechanism contributing to pain in DM1.

## Main Text

Myotonic dystrophy type 1 (DM1, Steinert’s disease) is the most prevalent form of adult muscular dystrophy. It arises from an unstable cytosine-thymine-guanine (CTG) trinucleotide repeat within the 3’ untranslated region of the protein kinase DM1 (*DMPK*) gene^1,2^. DM1 exhibits a multifaceted clinical presentation, encompassing muscle weakness, myotonia (delayed muscle relaxation), fatigue, cataracts, cardiac abnormalities, respiratory difficulties, gastrointestinal issues, cognitive changes, mood disturbances, and impaired fine motor skills^3^. Symptoms may vary in severity and can impact multiple organ systems, significantly impacting physical, physiological, and psychosocial aspects of afflicted persons’ lives^4–7^.

Chronic pain is a highly prevalent and often ignored DM1 symptom^8–13^. Pain is reported in more than 80% of DM1 patients with a significant correlation in pain severity and CTG nucleotide repeat size^8,12^. The pathophysiological origin of pain manifestation and maintenance in DM1 is completely unknown. Afflicted patients describe multiple qualities of pain, myalgia, and cramps that are widespread but most often located in the thighs, back, and proximal upper limbs. Many qualities of pain reported by DM1 patients (assessed via the short-form McGill pain questionnaire (SF-MPQ)) including “hot/burning,” “aching,” “heavy,” “tender,” and/or “tiring/exhausting” suggest a neuropathic origin for the pain phenotype^10^.

Despite the well characterized neuromuscular manifestations of DM1, the mechanisms responsible for pain in this disorder remain completely unknown. The high prevalence of pain, correlation with CTG repeat length, and frequent descriptions of burning, aching, and tenderness suggest that altered sensory processing may be a fundamental feature of the disease rather than simply a secondary consequence of muscle dysfunction. However, despite decades of work characterizing DM1 pathophysiology, pain has received remarkably little attention in preclinical studies. To our knowledge, pain-related behaviors and primary sensory neuron excitability have never been examined in an animal model of DM1. We recently demonstrated that both acute pharmacological and chronic genetic myotonia resulting from ClC-1 dysfunction produce pain-like behavior and alterations in nociceptive processing, suggesting that skeletal muscle hyperexcitability itself can engage sensory pain pathways^14^. Here, we asked whether a similar sensory phenotype is present in the context of DM1.

To address this question, we used the well-characterized HSA LR20b transgenic mouse model of DM1^15^. This genetically engineered mouse model carries a transgene for human skeletal actin (HSA) with an expanded CTG repeat sequence of ∼220 repeats in the 3’ untranslated region, mimicking the CTG repeat expansion in the *DMPK* gene seen in human DM1^15^. The expression of expanded CTG repeats triggers aberrant splicing of *Clcn1* in skeletal muscle and reduces the surface expression of voltage-gated chloride 1 (ClC-1) channels in muscle fibers producing recurrent myotonia in these mice^16,17^ **(Fig. 1A)**. Compared with wild-type littermates, DM1 mice exhibited significantly reduced withdrawal thresholds to punctate mechanical stimulation, increased responses to dynamic brush stimulation, exaggerated nocifensive behaviors following acetone application, and decreased withdrawal latencies on the hot plate test **(Fig. 1B-E)**.

**Figure 1.**
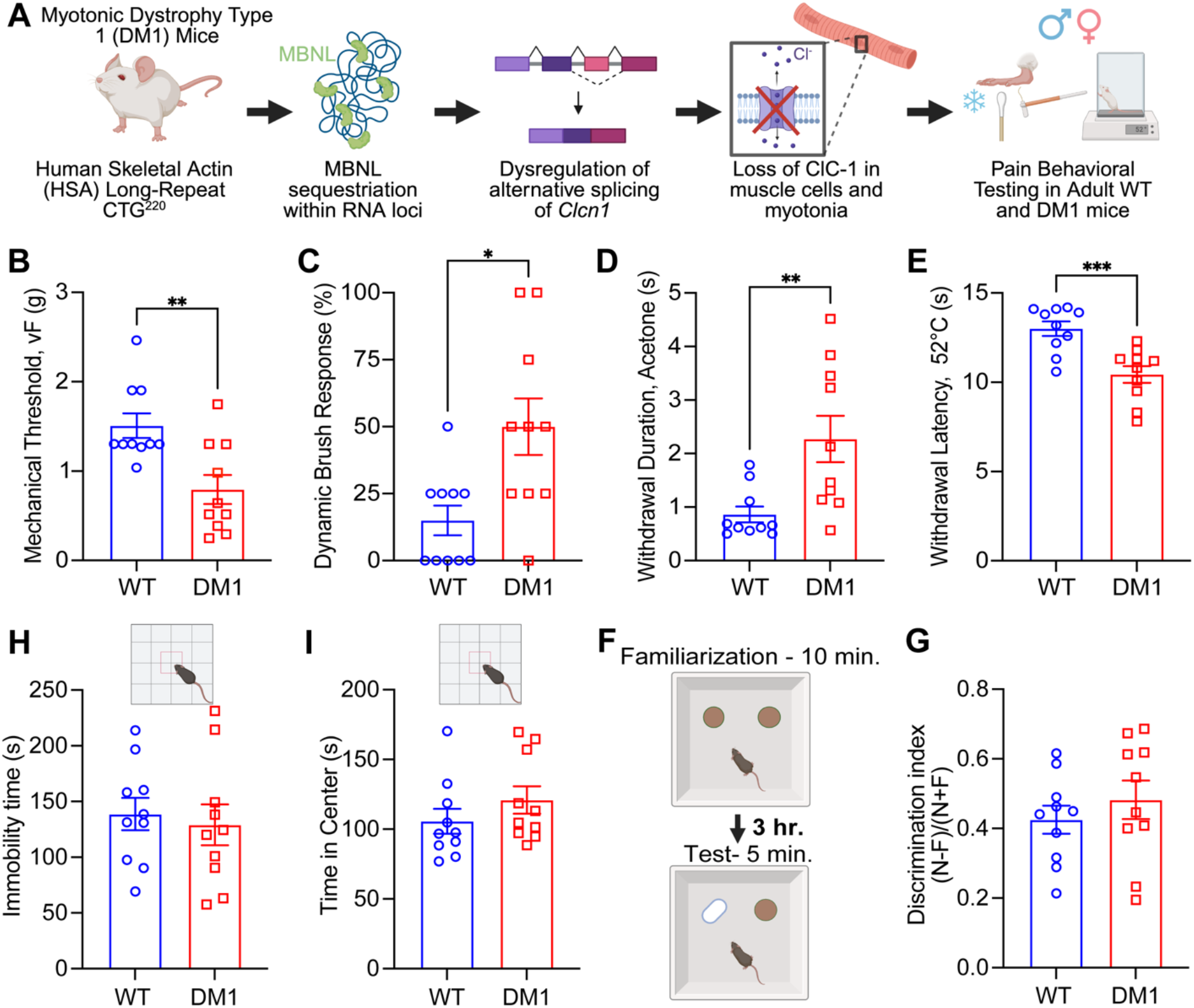
HSA LR20b myotonic dystrophy type 1 (DM1) mice exhibit multimodal hypersensitivity despite preserved locomotor activity and recognition memory. **(A)** Schematic illustrating the proposed mechanism by which expanded CTG repeats in the human skeletal actin (HSA) transgene of HSA LR20b mice result in muscleblind-like protein (MBNL) sequestration, aberrant pre-mRNA splicing, and dysregulation of *Clcn1*. Reduced ClC-1 chloride channel expression in skeletal muscle produces membrane hyperexcitability and myotonia. Adult wild-type (WT) FVB/NJ and HSA LR20b (DM1) mice were assessed for sensory and behavioral phenotypes. Compared with WT mice, DM1 mice exhibited significantly increased sensitivity to **(B)** punctate mechanical stimulation (von Frey), **(C)** dynamic mechanical stimulation (brush), **(D)** evaporative cooling (acetone), and **(E)** noxious heat (52°C hot plate). In contrast, DM1 mice exhibited normal **(F)** open-field locomotor activity and **(G)** novel object recognition performance. Data are presented as mean ± SEM (n = 10 mice/group). Individual data points represent values from single animals. Statistical analyses were performed using unpaired two-tailed Student’s *t* tests. *p < 0.05, **p < 0.01, ***p < 0.001.

Because DM1 is a multisystem disorder associated with cognitive impairment, fatigue, and motor deficits, we next determined whether the sensory phenotype occurred in the context of broader behavioral abnormalities. However, DM1 mice exhibited normal locomotor activity in the open field and intact novel object recognition performance **(Fig. 1F,G)**. These findings indicate that DM1 mice exhibit robust multimodal hypersensitivity despite preserved locomotor and cognitive function, suggesting that altered somatosensory processing is a selective feature of the disease rather than a manifestation of generalized neurological dysfunction.

Because chronic pain states are frequently associated with increased excitability of primary sensory neurons, we next examined the intrinsic electrophysiological properties of dorsal root ganglion (DRG) neurons isolated from adult DM1 and wild-type mice **(Fig. 2A)**. Representative traces revealed increased action potential firing in small- and medium-diameter sensory neurons from DM1 mice following ramp current injection **(Fig. 2B)**. Quantification demonstrated a significant depolarization of resting membrane potential and an increase in evoked action potential firing, whereas rheobase was unchanged **(Fig. 2C-E)**. In contrast, large-diameter sensory neurons exhibited no differences in resting membrane potential or rheobase between genotypes **(Fig. 2F,G)**. Together, these findings demonstrate that DM1 mice exhibit robust multimodal hypersensitivity accompanied by selective hyperexcitability of putative nociceptive sensory neurons, identifying peripheral sensitization as a potential contributor to pain in DM1.

**Figure 2.**
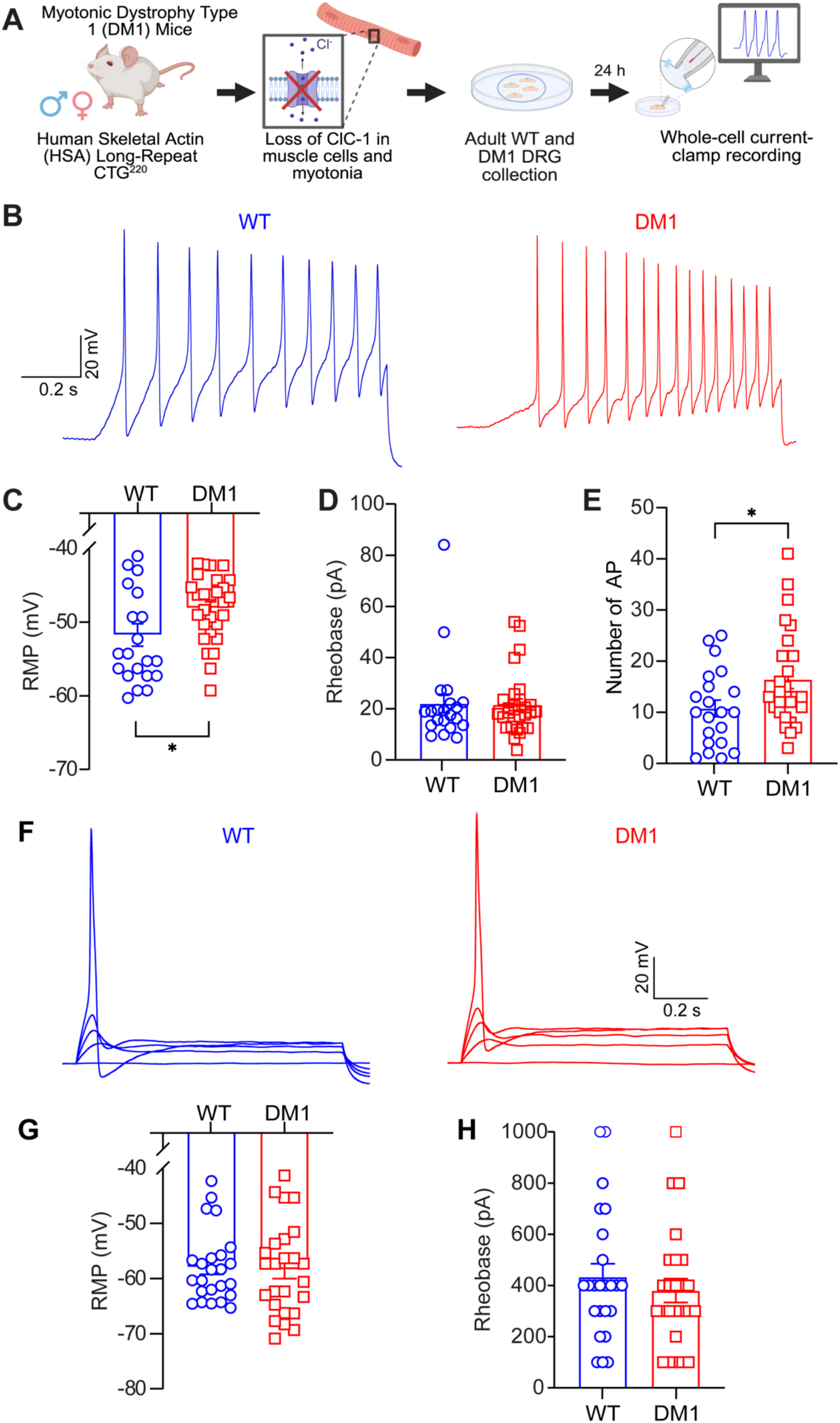
HSA LR20b myotonic dystrophy type 1 (DM1) mice exhibit selective hyperexcitability of putative nociceptive dorsal root ganglion neurons. **(A)** Schematic illustrating the experimental workflow. Lumbar dorsal root ganglia (DRGs) were isolated from adult wild-type (WT) and HSA LR20b (DM1) mice and dissociated for whole-cell current-clamp electrophysiological recordings. **(B)** Representative traces showing action potential (AP) firing in small- and medium-diameter sensory neurons from WT (left) and DM1 (right) mice in response to depolarizing ramp current injection. Quantification of **(C)** resting membrane potential, **(D)** rheobase, and **(E)** number of evoked APs demonstrated increased excitability of small- and medium-diameter neurons from DM1 mice. In contrast, large-diameter sensory neurons exhibited no differences in **(F)** resting membrane potential or (G) rheobase between genotypes. Data are presented as mean ± SEM. Individual data points represent single neurons. Small- and medium-diameter neuron recordings were obtained from n = 21–28 neurons from 4 mice/group. Large-diameter neuron recordings were obtained from n = 24–25 neurons from 4 mice/group. Statistical analyses were performed using Mann-Whitney U tests. *p < 0.05.

The present findings identify a previously unrecognized sensory phenotype in DM1 mice characterized by multimodal hypersensitivity and selective hyperexcitability of putative nociceptive DRG neurons. Importantly, these abnormalities occurred despite preserved locomotor activity and recognition memory, suggesting that expanded CTG repeats are associated with alterations in somatosensory processing rather than generalized behavioral dysfunction.

This preclinical phenotype is broadly consistent with the emerging clinical picture of pain in DM1. Pain affects the majority of patients with DM1 and is increasingly recognized as a major determinant of impaired daily functioning and reduced quality of life^8,9,12,18^. Moreover, pain severity correlates with disease duration, CTG repeat size, fatigue, and other measures of disease burden^8,12^. Recent sensory phenotyping studies have further demonstrated reduced thermal and mechanical pain thresholds, pressure hyperalgesia, and evidence of small-fiber pathology in subsets of patients with DM1^19–22^. Together, these observations support the concept that pain in DM1 reflects, at least in part, abnormalities in sensory processing rather than simply a secondary consequence of muscle weakness or disability.

The present electrophysiological findings provide a potential mechanistic basis for these clinical observations. Small- and medium-diameter DRG neurons from DM1 mice exhibited depolarized resting membrane potentials and increased action potential firing, findings indicative of enhanced neuronal excitability and consistent with peripheral sensitization. Depolarized membrane potentials could result from reduced potassium conductance, such as diminished activity of leak channels or inward rectifiers^23^, but this was not directly tested. Additionally, while DM1 DRGs showed increased firing, the specific ion channels contributing to this hyperexcitability remain undefined. Likely candidates include voltage-gated sodium (NaV) channels, which play central roles in shaping neuronal firing thresholds and frequency^24^. Although the mechanisms responsible for this hyperexcitability remain undefined, our recent work demonstrated that transient pharmacological and chronic genetic myotonia are sufficient to produce persistent pain-like behavior and altered sensory neuron excitability^14^. These findings raise the possibility that persistent myotonia and repetitive muscle contractions in DM1 provide sustained nociceptive input to muscle-innervating afferents and induce activity-dependent plasticity within the peripheral sensory nervous system. Future electrophysiological studies using voltage-clamp techniques and pharmacological tools will be essential to dissect the individual channel contributions and fully elucidate the mechanisms driving sensory neuron sensitization in chronic myotonic disease.

Several limitations warrant consideration. The HSA LR20b model primarily recapitulates the muscle pathology and myotonia associated with DM1 and therefore may not fully capture CTG repeat-associated abnormalities in other organ systems, including the nervous system. Consequently, additional mechanisms may contribute to pain manifestation and maintenance in patients with DM1. Furthermore, although our findings are consistent with peripheral sensitization, we did not directly assess alterations in central nociceptive circuits and therefore cannot exclude contributions from central sensitization. Finally, the molecular and cellular mechanisms responsible for DRG hyperexcitability remain undefined, and future studies will be required to determine the ion channels, signaling pathways, and repeat-associated processes linking chronic myotonia to sensory neuron dysfunction.

Nevertheless, our findings establish HSA LR20b mice as a translational model for investigating pain in DM1 and provide the first preclinical evidence that DM1 is associated with multimodal hypersensitivity and selective hyperexcitability of primary sensory neurons. Together with our recent demonstration that myotonic muscle activity is sufficient to initiate persistent alterations in nociceptive processing^14,25^, these findings support an emerging framework in which pathological skeletal muscle activity contributes directly to sensory nervous system dysfunction and pain across myotonic disorders. Together, these findings support the concept that altered sensory processing is a clinically relevant feature of DM1 and identify peripheral sensory neuron dysfunction as a plausible contributor to this common and disabling symptom.

## Methods

### Animals

Adult FVB/NJ (Jackson Laboratory, #001800) and FVB/N-Tg(HSA*LR)20bCath/J (Jackson Laboratory, #032031) male and female mice were group housed under temperature- and humidity-controlled conditions and maintained on a 12:12 hour light cycle with food and water available ad libitum. Adult mice between 9 weeks and 3 months of age were used for all experiments. Equal numbers of male and female mice were used for all tests. Although the study was not powered to detect sex differences, no obvious trends were observed and data from both sexes were pooled for analysis. All experimental procedures were approved by the Institutional Animal Care and Use Committees of University of Florida (IACUC202400000002) and New York University (PROTO202100104) and were performed in accordance with the National Institutes of Health Guide for the Care and Use of Laboratory Animals.

### Behavioral Testing

Mice were acclimated to the behavioral testing room for at least 30 minutes prior to experimentation. Experimenters were blinded to genotype throughout all behavioral assessments.

#### Static mechanical sensitivity (von Frey)

As previously described^26,27^, mice were habituated for 60 minutes in plexiglass chambers positioned on an elevated wire mesh platform. Calibrated von Frey filaments (0.007–6.0 g; Braintree Scientific) were applied to the lateral plantar surface of the hindpaw using the up-down method, and the 50% withdrawal threshold was calculated.

#### Dynamic mechanical sensitivity (brush)

As previously described^26^, immediately following von Frey testing, the plantar surface of the hindpaw was gently brushed in a heel-to-toe direction using a puffed cotton swab. Four stimulations were performed and the frequency of withdrawal responses was recorded.

#### Cold sensitivity (acetone)

As previously described^26,27^, approximately 10 μL of acetone was applied to the plantar hindpaw using a syringe connected to flared PE-90 tubing. The cumulative duration of paw lifting, licking, or shaking was recorded during a 30-second observation period. Three trials were averaged for each animal.

#### Thermal sensitivity (hot plate)

As previously described^26,28^, mice were placed on a 52.5°C hot plate (Ugo Basile), and the latency to a nociceptive response, defined as paw licking or jumping, was recorded. A cutoff time of 30 seconds was imposed to avoid tissue injury. One to three trials separated by at least 10 minutes were averaged for each animal.

#### Open field Test (OFT)

Open-field testing was performed to assess spontaneous locomotor activity and exploratory behavior. Mice were placed individually into a square open-field arena (30 × 30 cm) and allowed to freely explore for 15 minutes. Behavior was recorded using an overhead camera (Microsoft LifeCam HD-3000) and analyzed using ANY-maze software (version 7.2, Stoelting Co). Total distance traveled was quantified and used as a measure of locomotor activity. Upon completion of the open-field session, mice were removed from the arena, and the same apparatus was subsequently used for novel object recognition testing on the following day, with the open-field session serving as habituation to the testing environment.

#### Novel object recognition (NOR

Novel object recognition testing was performed in the same arena as the open-field test, allowing the open-field session to serve as habituation to the testing environment. A camera positioned above the arena was connected to ANY-maze software (Stoelting) for live tracking and video recording. Two identical objects were placed in opposite quadrants of the arena before the mice were reintroduced to the arena and allowed to freely explore the objects for 10 minutes (familiarization phase). After a 3-hour retention interval, one familiar object was replaced with a novel object and mice were returned to the arena for a 5-minute test session. The location of the novel object was counterbalanced between mice to minimize potential side preferences. Exploration was defined as the mouse directing its nose toward an object within approximately 2 cm while actively investigating it. Recognition memory was assessed using the discrimination index calculated as:

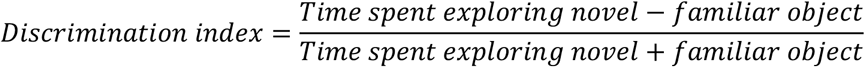

Positive discrimination index values indicate a preference for the novel object and intact recognition memory.

### Electrophysiological Recordings in Dorsal Root Ganglion Neurons

Whole-cell patch-clamp recordings were performed in dissociated lumbar dorsal root ganglion (DRG) neurons using procedures previously established by our laboratory^14,29^. Lumbar DRGs were harvested from adult male and female mice and enzymatically dissociated for 50 minutes at 37°C in Dulbecco’s Modified Eagle Medium containing collagenase type I (1.66 mg/mL; Worthington) and neutral protease (1.04 mg/mL; Worthington). Dissociated neurons were plated onto poly-D-lysine-coated coverslips and maintained in culture medium supplemented with 10% fetal bovine serum and 1% penicillin/streptomycin. Recordings were performed 15–24 hours after plating.

Whole-cell current-clamp recordings were acquired at room temperature (20–22°C) using an EPC 10 amplifier (HEKA Elektronik, Germany) controlled by PatchMaster software. Borosilicate glass pipettes (2–3.5 MΩ) were filled with intracellular solution. Experimenters were blinded to genotype throughout data acquisition and analysis.

Neurons were categorized according to cell size and whole-cell capacitance. Small- and medium-diameter neurons were defined as having capacitance values less than 15 pF and approximate diameters less than 23 μm. Large-diameter neurons exhibited capacitance values greater than 40 pF and approximate diameters of 37 μm. In small- and medium-diameter neurons, action potentials were elicited using a depolarizing ramp protocol (0–150 or 0–250 pA over 1 second), and rheobase was defined as the minimal current required to evoke an action potential. Because large-diameter neurons were not amenable to ramp protocols, rheobase was determined using 1-second depolarizing current injections delivered in 100 pA increments.

### Blinding and Randomization

Experimenters were blinded to genotype during all behavioral and electrophysiological experiments. Animals were assigned to experimental groups according to genotype, and data analysis was completed before group identities were revealed.

### Statistics

All data are presented as mean ± SEM. Individual data points represent single animals for behavioral experiments and individual neurons for electrophysiological recordings. Statistical analyses were performed using GraphPad Prism (v10.2.3). Comparisons between two groups were performed using unpaired two-tailed Student’s t tests or Mann-Whitney U tests, as appropriate following assessment of data distribution. Exact sample sizes, statistical tests, and p values are reported in the figure legends. Statistical significance was defined as p < 0.05. No formal power calculations were performed, and sample sizes were based on prior experience with similar behavioral and electrophysiological assays.

## Data Availability

All data is available by the corresponding author on reasonable request.

## Code Availability

No custom code was generated.

## Acknowledgements

This work was supported was supported by National Institutes of Health (NIH) awards K00NS124190 (to TSN), K99NS134965 (to KG), and RF1NS131165, R61NS126026, R01NS120663 (to RK). Additionally, this work was supported by a Development Grant from the American Neuromuscular Foundation (to TSN).

## Author contributions

TSN conceived the project, designed and performed experiments, and wrote the manuscript. ACR led the DRG electrophysiological recordings in dorsal root ganglion neurons. SH, SLL, KG, and PD contributed to data acquisition. RK directed the overall project, secured funding, and provided scientific oversight. All authors contributed to manuscript editing and approved the final version.

